# Mice born preterm develop gait dystonia and reduced cortical parvalbumin immunoreactivity

**DOI:** 10.1101/2024.02.01.578353

**Authors:** Kat Gemperli, Femi Folorunso, Benjamin Norin, Rebecca Joshua, Clayton Hill, Rachel Rykowski, Rafael Galindo, Bhooma R. Aravamuthan

## Abstract

Preterm birth leading to cerebral palsy (CP) is the most common cause of childhood dystonia, a movement disorder that is debilitating and often treatment refractory. Dystonia has been typically associated with dysfunction of striatal cholinergic interneurons, but clinical imaging data suggests that cortical injury may best predict dystonia following preterm birth. Furthermore, abnormal sensorimotor cortex inhibition has been found in many studies of non-CP dystonias. To assess the potential for a cortical etiology of dystonia following preterm birth, we developed a new model of preterm birth in mice. Noting that term delivery in mice on a C57BL/6J background is embryonic day 19.1 (E19.1), we induced preterm birth at the limits of pup viability at embryonic day (E) 18.3, equivalent to human 22 weeks gestation. Mice born preterm demonstrate display clinically validated metrics of dystonia during gait (leg adduction amplitude and variability) and also demonstrate reduced parvalbumin immunoreactivity in the sensorimotor cortex, suggesting dysfunction of cortical parvalbumin-positive inhibitory interneurons. Notably, reduced parvalbumin immunoreactivity or changes in parvalbumin-positive neuronal number were not observed in the striatum. These data support the association between cortical dysfunction and dystonia following preterm birth. We propose that our mouse model of preterm birth can be used to study this association and potentially also study other sequelae of extreme prematurity.

## Introduction

In the United States, preterm birth is the most common cause of cerebral palsy (CP), a lifelong motor disability affecting between 2-4 of every 1000 people.(Boyle et al., 2011; McIntyre et al., 2022; Van Naarden Braun et al., 2016) As a result, preterm birth is also the most common cause of dystonia, which can affect up to 80% of people with CP.(Rice et al., 2017) Dystonia is formally defined as voluntary-movement triggered overflow muscle activation and can be worsened with fatiguing or increasingly challenging voluntary movement tasks, making dystonia an inherently disabling movement disorder.(Albanese et al., 2013; Sanger et al., 2010, 2003) Dystonia is notoriously difficult to diagnose in people with CP and is often under-diagnosed, precluding treatment.(Bhooma R Aravamuthan et al., 2023; Miao et al., 2022) Furthermore, commonly used treatments, like anticholinergic agents, are variably effective at treating dystonia in CP.(Bohn et al., 2021) This necessitates dedicated investigation of treatment targets specific to dystonia following preterm birth.

Identification of dystonia treatment targets specific to preterm birth requires a clinically relevant model of preterm birth and clinically relevant methods to quantify dystonia in mice. Mouse models of prematurity have been difficult to develop because of differences in the mouse neurodevelopmental timeline and human neurodevelopmental timeline.(Clancy et al., 2007; Semple et al., 2013) Based on rate of brain growth and myelination patterns, postnatal day 10 (P10) in the mouse is thought to be equivalent to human term gestation. The day of birth in the mouse (P0) is thought to be equivalent to human 24 weeks gestation.(Clancy et al., 2007; Semple et al., 2013) Existing mouse models of prematurity involve hypoxic or inflammatory insults prior to putative human equivalent gestation (P10) but do not result in true preterm birth of the mouse.(Le Dieu-Lugon et al., 2020; McCarthy et al., 2018) Existing mouse models of true preterm birth focus on outcomes to the dam without assessment of outcomes in viable pups.(McCarthy et al., 2018) Examining the outcomes of true preterm birth in pups is additionally important because lower gestational ages contribute to the highest morbidity. For example, a child born between 22-24 weeks gestation has almost double the risk of CP compared to a child born between 25-27 weeks gestation.(Boyle et al., 2011; Chen et al., 2021; Van Naarden Braun et al., 2016) Therefore, an ideal model of mouse prematurity would replicate true preterm birth.

Assessing dystonia in mouse models of disease is difficult in large part because clinical diagnosis is difficult. Gold standard clinical diagnosis involves qualitative consensus-based expert review of neurologic exam videos, which is obviously a difficult practice to translate to animal models. Identification of dystonia in mice has canonically involved identification of clasping during tail suspension, a phenomenon where mice adduct their limbs to midline and clasp their paws together.(Oleas et al., 2013; Richter and Richter, 2014; Tassone et al., 2011) Unfortunately, this phenomenon is observed primarily during tail suspension, a task that is difficult to assess in people, and is not present in more clinically relevant tasks like gait. Furthermore, the typical assessment of limb adduction culminating in clasping is binary (i.e. clasping is present or absent), precluding quantification of more subtle limb adduction that may still be dystonic but is not formally clasping. We have demonstrated that limb adduction amplitude and variability during gait is cited by experts when differentiating dystonia from spasticity and when assessing dystonia severity in people with CP.(Aravamuthan et al., 2021; Bhooma R. Aravamuthan et al., 2023) We have also demonstrated quantifiable metrics of leg adduction amplitude and variability track with expert assessments of dystonia severity in people with CP.(Bhooma R. Aravamuthan et al., 2023; Gemperli et al., 2023) Therefore, we propose that quantifying leg adduction amplitude and variability during gait in mice is a clinically-relevant metric of dystonia.

Dystonia is typically associated with striatal injury, particularly striatal cholinergic interneuron (ChIN) excitation which has been viewed as a unifying mechanism of non-CP dystonia etiologies like DYT1 and DYT6.(Bonsi et al., 2011; Chintalapati et al., 2020; Eskow Jaunarajs et al., 2019, 2015; Gemperli et al., 2023; Pappas et al., 2015) By measuring leg adduction variability and amplitude, we have also shown that chronic striatal ChIN excitation can cause dystonia.(Gemperli et al., 2023) Finally, we have shown that a rat model of hypoxic-ischemic injury at human-equivalent *term* gestation demonstrates an increased number of striatal ChINs.(Gandham et al., 2020) However, despite this evidence implicating abnormal striatal ChIN excitation in dystonia pathogenesis, anticholinergic agents are often ineffective to treat dystonia in CP.(Bohn et al., 2021) This suggests involvement of other brain regions in dystonia pathogenesis. We have shown that cortical injury may cause dystonia in people born preterm.(Chintalapati et al., 2023; Ueda et al., 2023) Over half of people with CP with cortical injury alone on MRI have dystonia.(Ueda et al., 2023) Furthermore, a marker of cortical atrophy, more so than striatal volume loss, best correlates with the presence of dystonia in people with CP born preterm.(Chintalapati et al., 2023) Lesion mapping,(Corp et al., 2019) PET,(Garibotto et al., 2011) electrocorticography,(Miocinovic et al., 2015) and fMRI data(Norris et al., 2020) all also implicate abnormal inhibition(Gallea et al., 2017) of the sensorimotor cortex in the pathogenesis of non-CP dystonia, suggesting involvement of cortical inhibitory interneurons. The largest population of inhibitory neurons in the cortex are parvalbumin-positive interneurons (PVINs).(Lim et al., 2018; Nahar et al., 2021; Tremblay et al., 2016) Cortical PVINs are lost or have reduced activity in many mouse models of autism(Juarez and Martínez Cerdeño, 2022) and epilepsy,(Jiang et al., 2016) symptoms that commonly occur in people with CP.(Dos Santos Rufino et al., 2023; Påhlman et al., 2021) However, it is unclear if they are involved in dystonia pathogenesis following preterm birth.

Here, we demonstrate that dystonia, as measured by the clinically-relevant metrics of leg adduction variability and adduction, is present in our novel mouse model of preterm birth and extreme prematurity. We also show that this model has decreased parvalbumin immunoreactivity that is restricted to the cortex and is not present in the striatum. We propose that this model can be used to study the cortical pathophysiology of dystonia following preterm birth.

## Methods

All procedures were approved by the Washington University in St. Louis Institutional Animal Care and Use Committee (protocol # 21-0174) and were in strict accordance with the National Institute of Health Guidelines on the Care and Use of Laboratory Animals. All mice used for these experiments were on a C57BL/6J background and obtained from Jackson Labs. Mice were housed in a reverse light cycle facility (lights on from 6PM to 6AM).

### Breeding and induction of labor

Mice were bred between 4-6 PM on alternate days and checked for mucus plugs at 6 PM to determine the time of conception. The documentation of a mucus plug was timed as embryonic day 0 and mice were weighed on that day to establish their pre-pregnancy weight. To induce labor, mice were subcutaneously injected with mifepristone (or vehicle for controls) at E17.6 (8 AM on 18^th^ day after the plug was noted). Of note, though mifepristone is known to cross the placenta, there is no evidence for adverse neurologic effects from mifepristone based on human data or from data examining mouse pup outcomes or cortical injury patterns when delivered at term gestation (E19.5).(Bernard et al., 2013; HILL et al., 1990; Morin et al., 2022) Mifepristone (Sigma Aldrich, M8046) was dosed at 3 μg per gram of pre-pregnancy weight using a 1 µg/µL, suspended in 100% ethanol. Dams delivering term-born controls were given the weight-based dosing of an equivalent volume of ethanol. Mifepristone injections were timed to induce delivery at E18.3, 18 hours after injection, at human equivalent 22 weeks gestation. This gestational age was chosen to limit pup mortality noting that pup skin integrity (impermeability to extraneously applied dyes) develops at E18.

All deliveries were filmed to calculate gestational age at birth using a GardePro E7 WiFi Trail Camera. The time of birth was documented as the 5 minute window in which the first pup was seen as delivered (postnatal day 0, or P0). Gestational age was calculated as P0-E0.

### Motor behavioral assessment

Mouse gait was assessed in adolescence at P42, after gait patterns are fully mature. We assessed gait in two ways: via treadmill gait and via open field. We constructed a clear treadmill using ¼” thick acrylic sheets with a walkway 1.5” in diameter and 10.5” in length that was elevated 11.5” off the ground (Figure 1). A rotary motor (Tsiny 2GN-90K 110V DC Gear-Box) was used to run an acetate sheet (4mm thickness) over the treadmill. Mice were placed on this treadmill belt and were filmed while walking for 30 seconds each at 3 different speeds: 5, 8, and 11 cm/second.

**Figure 1.**
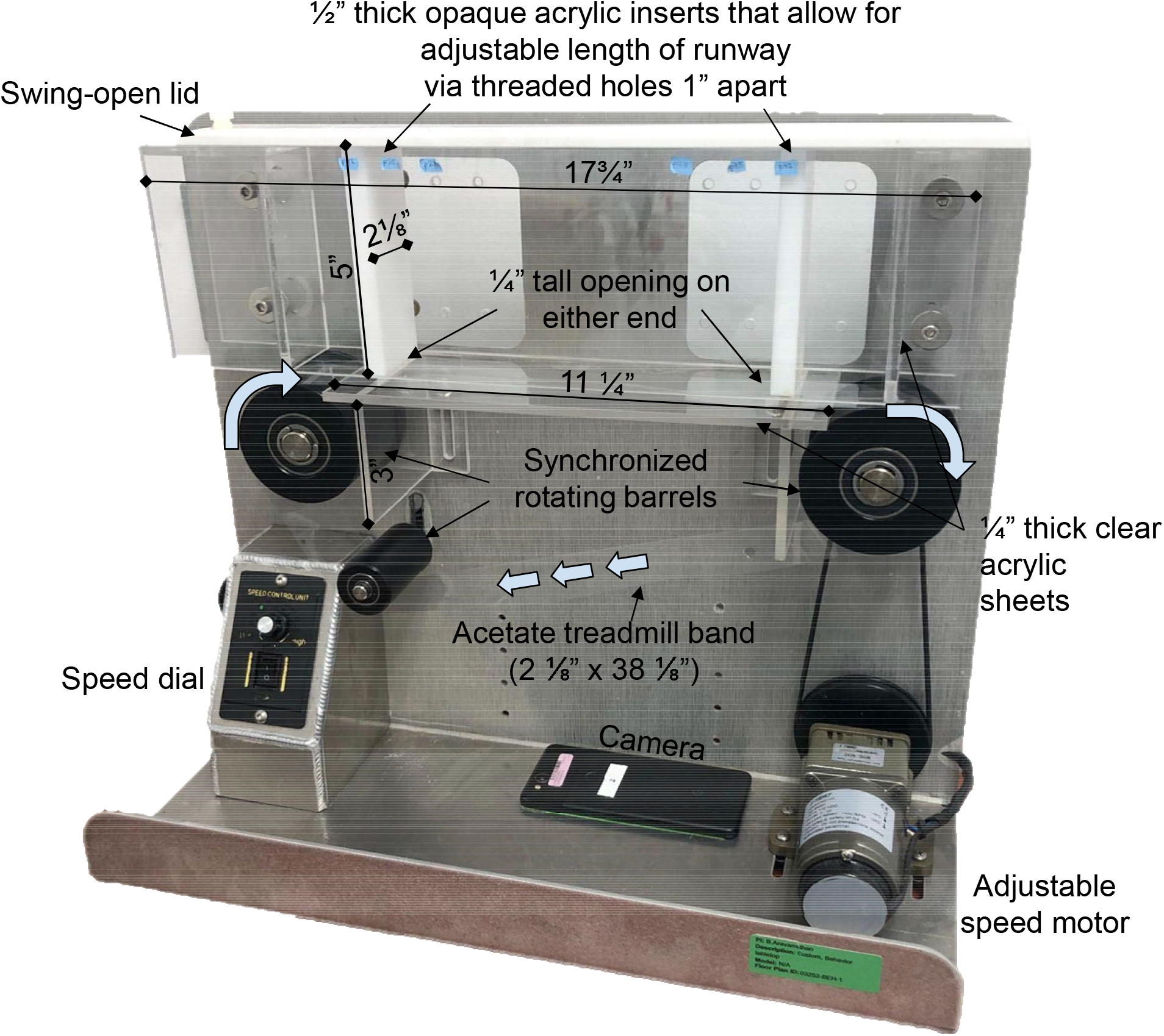
Treadmill design.

Open field gait was assessed by placing mice in a 50cm x 50cm clear acrylic box placed on a clear acrylic top table 3 feet tall. Walls were obscured with blue film, but the bottom of the box remained transparent. This allowed for filming from underneath the mouse. Mice were placed in the center of the box and the top of the box was covered with another blue sheet. Spontaneous mouse movements were recorded for 10 minutes.

Videos for both treadmill and open field. were recorded from underneath each mouse such that the ventral surface of the mouse and foot placement was visible on camera. Videos were recorded using a Google Pixel 2 (Google, Mountain View, CA, USA) at 120 frames per second at 1980 × 1020 pixel resolution.

### Quantification of locomotor impairment and dystonia

We used open source pose estimation software called DeepLabCut(Mathis et al., 2018; Nath et al., 2019) (https://github.com/DeepLabCut/) to train neural network models to label the following points of interest on each mouse in each recorded video: nose, body midpoint, tail base, left and right fore- and hindpaw tops (middle toes) and bases (pad of each paw). For open field, the four corners of the open field box were also labeled. To train these models, twenty frames were extracted from each video used for training (42 for treadmill, 28 for open field) using k-means extraction. We labeled points of interest manually on these frames and these labels were used to train and test separate models to label the above points on mice across all videos recorded for each task. Models were ResNet-50-based neural network trained for at least 500,000 iterations with 95% of frames used for training and 5% of frames used for testing the network to yield less than a 5 pixel train error and 5 pixel test error. These models generate X and Y coordinates for each labeled point and also a p-value indicative of the likelihood of correct labeling. If a point is not visible on a frame, its p-value is low (minimum 0). If a point is easily identifiable on a frame, its p-value is high (maximum 1). All coordinates used for analysis had a p-value greater than 0.99.

Locomotion was assessed using the following metrics:(Broom et al., 2017)

- For both open field and treadmill gait: number of steps taken and percent time spent in bipedal support vs. tripedal/quadrupedal support (analogous to human single support vs. double support). We note that children with CP spend a decreased amount of time in the single support gait phase and, consequently, spend increased time in the double support gait phase to compensate for decreased gait stability.(Brégou Bourgeois et al., 2014; Carcreff et al., 2020) The analogous finding in mice would be decreased time spent in bipedal support.
- For open field alone: total distance traveled (cm) and maximum speed during a single second epoch (cm/sec) were also calculated (which could not be calculated for treadmill due to the set distance and speeds traveled).

Dystonia was assessed using hindlimb foot angles (the angle between each hindpaw top, base, and tailbase). Angles were only calculated when the mouse was walking in a straight line (nose to body midpoint to tail base angle between 170-190 degrees) and was walking at a speed of at least 5 cm/sec (either on treadmill or open field) to ensure a consistent gait pattern when assessing foot angles. Foot angle minimum and foot angle variance were used to measure limb adduction amplitude and variability, as we have previously demonstrated.(Bhooma R. Aravamuthan et al., 2023; Gemperli et al., 2023) All calculations were done using custom written code in MATLAB.

Trained DeepLabCut models and MATLAB analysis code are available here: https://wustl.box.com/s/xzshevre9zsp0z6xi5qyfkzn6stfycn0

### Brain tissue processing and immunohistochemistry

Mice were anesthetized deeply with isoflurane and transcardially perfused with phosphate-buffered saline (PBS) and then 4% paraformaldehyde diluted in PBS. Brains were extracted and then stored in 4% paraformaldehyde for 24 hours before cryoprotection with 30% sucrose in PBS for at least 24 hours before further processing. Brains were sliced in 50 μ m sections using a freezing microtome. Two slices per mouse were used for quantification at 1.1 mm and 0.7 mm anterior to bregma, regions encompassing both the sensorimotor cortex (defined as the adjacent primary motor and primary sensory cortices) and the striatum. To evaluate psarvalbumin immunoreactivity, free floating slices were immunohistochemically stained for parvalbumin (Primary: anti-Parvalbumin, MilliporeSigma MAB1572, 1:500 dilution; Secondary: goat anti-mouse Alexa Fluor^™^ 568, ThermoFisher Scientific, 1:500 dilution). Brain sections were then mounted in Fluoromount (ThermoFisher Scientific, Waltham, MA) and scanned using a Hamamatsu NanoZoomer 2.0 digital slide scanner (Hamamatsu Photonics K.K., Shizuoka, Japan).

### Parvalbumin immunoreactivity quantification

Images were calibrated for identical brightness levels (68%) post-imaging using NDP.view2 software (Hamamatsu Photonics K.K., Shizuoka, Japan). This brightness allowed for optimal viewing of neuronal staining without contamination from background staining across all brain slices. Images were further analyzed using ImageJ. Regions of interest (ROIs) were drawn bilaterally on a single non-experimental slice at 0.7 mm anterior to Bregma. These ROIs encompassed: 1) the primary motor and primary sensory cortices using the corpus callosum as a landmark, 2) the dorsal striatum using the corpus callosum as the superior and lateral borders, the lateral ventricle as the medial border, and the anterior commissure as the inferior border). The same ROIs were used on to quantify parvalbumin immunoreactivity on every slice assessed to ensure comparable areas were assessed for each brain. Slices were set to 8-bit grayscale and then thresholded identically using the MaxEntropy profile with a range of 39-255. ROIs were then overlayed on these thresholded slices. For quantifying parvalbumin immunoreactivity in the sensorimotor cortex, we calculated the percent area occupied by the thresholded signal within each sensorimotor cortex ROI. For quantifying parvalbumin immunoreactivity in the striatum, individual neuron bodies were manually counted for each ROI for each slice. Notably, throughout all quantification, the experimenter was blinded to the cage ID and experimental group of each mouse. To further avoid bias from a single slice or ROI, all values generated for each brain region were averaged to generate a single value for parvalbumin immunoreactivity for the sensorimotor cortex and a single value for parvalbumin neuron number for the striatum for each mouse.

### Statistics

All statistical analysis was done in GraphPad Prism (version 8, GraphPad Software). Parvalbumin immunoreactivity, locomotor impairment metrics during open field, and dystonia metrics during open field were compared between term and preterm mice using t-tests when appropriate (with normality of data sets assessed using the Shapiro-Wilk test). For data with unequal standard deviations between groups, a t-test with Welch’s correction was applied. For data that were not normally distributed, Mann-Whitney tests were used. Locomotor impairment metrics and dystonia metrics during treadmill gait were compared between term and preterm mice across three different treadmill belt speeds using a two-way repeated measures ANOVA.. The significance level for all tests was set a priori to p<0.05.

### Data availability

The authors confirm that the data supporting the findings of this study are available from qualified investigators upon request.

## Results

We generated 5 litters born at term gestation (mean gestational age 19.12 days, 95% CI 19.05-19.19, range 19.03-19.23) yielding 34 term-born pups and 6 litters born at preterm gestation (mean gestational age 18.31 days, 95% CI 18.28-18.35, range 18.23-18.38) yielding 29 pups born preterm. Litter sizes trended toward being smaller for preterm born mice (3-6 pups per litter) compared to term born mice (5-9 pups per litter) but were not significantly different between groups (Figure 2).

**Figure 2.**
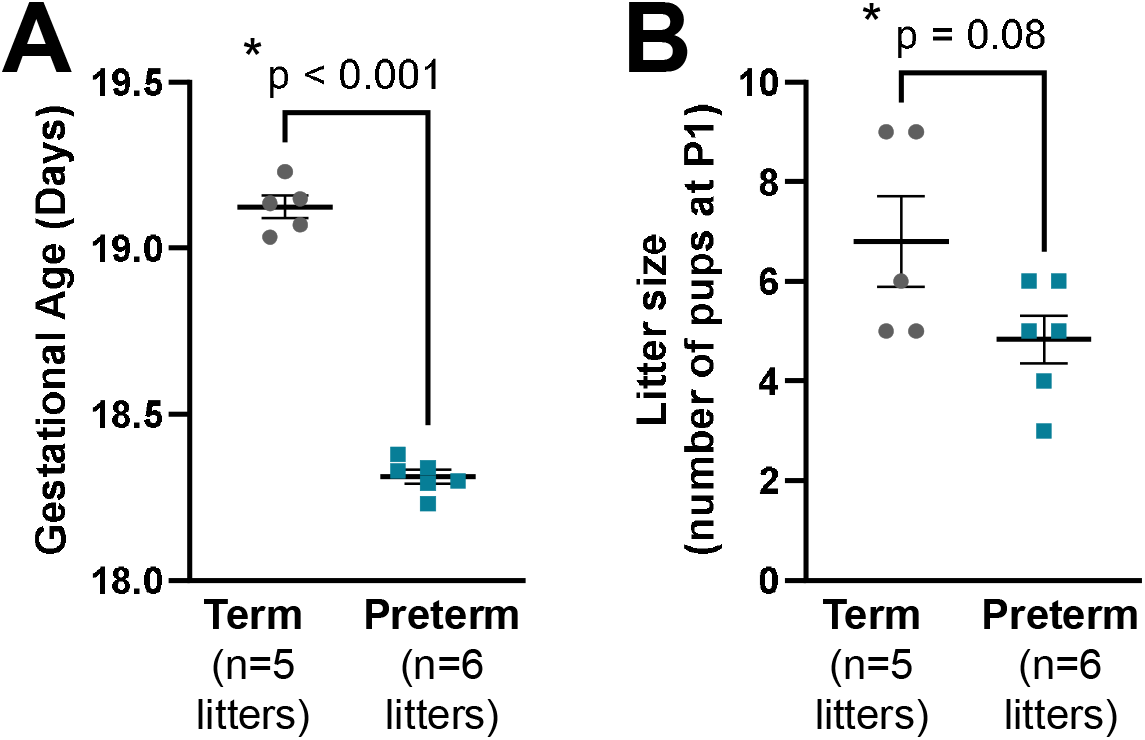
Gestational ages (A) and litter sizes (B) of mice born preterm (n=6 litters) and term-born controls (n=5 litters). *T-test

Mice born preterm demonstrated subtle, but nonetheless clinically relevant, motor impairments during treadmill and open field gait. During both tasks, mice born preterm demonstrated significantly reduced time spent in bipedal support compared to term born controls, suggesting diminished gait stability requiring increased duration tripedal or quadrupedal gait. There was no difference in number of steps taken during either task or in total distance traveled or maximum speed during the open field task (Figure 3).

**Figure 3.**
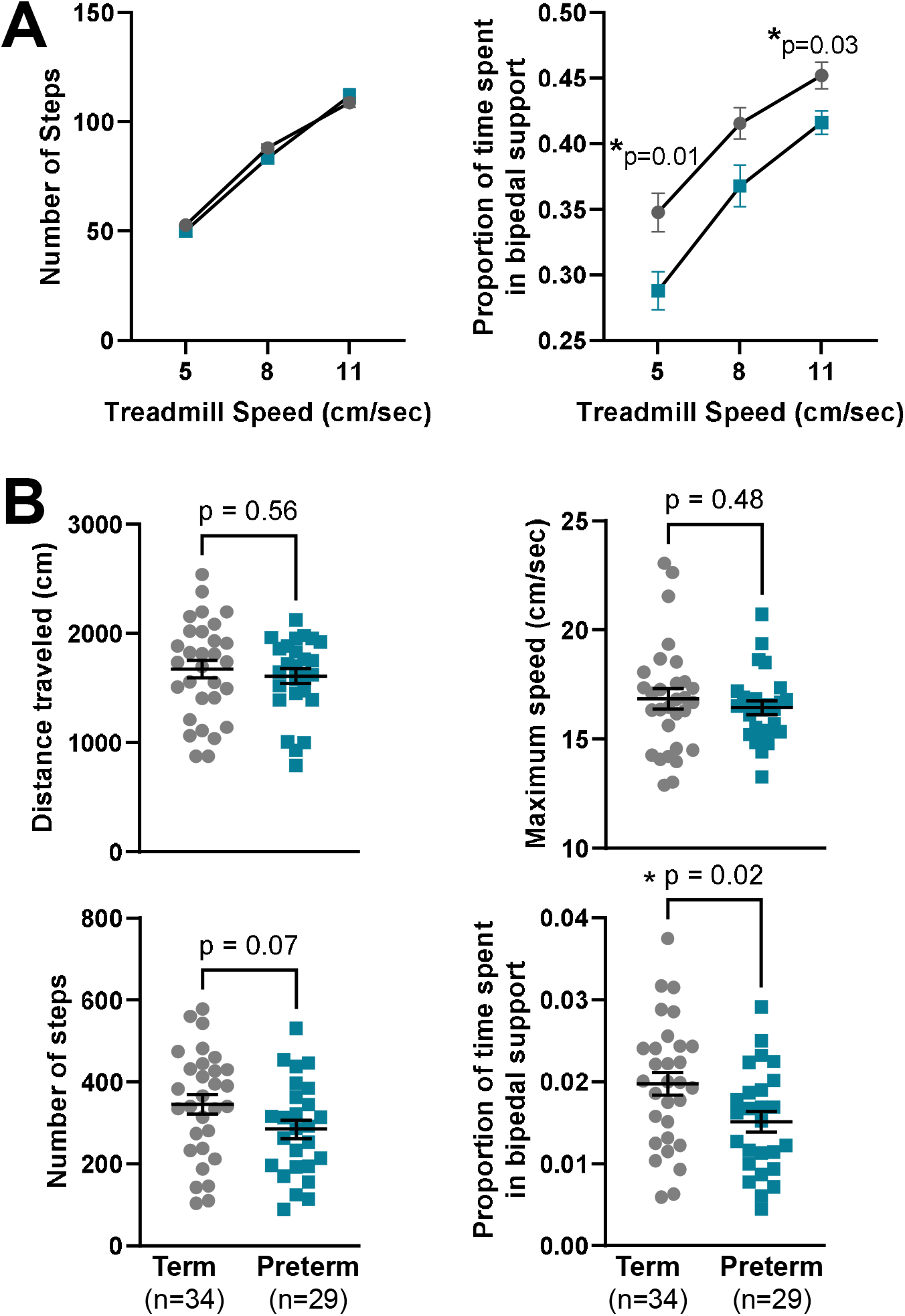
Locomotor impairment during treadmill gait (A, two-way repeated measures ANOVA) and Open field (B, T-test). Mice born preterm demonstrate a reduced proportion of time in bipedal support during gait.

Mice born preterm also demonstrated dystonic gait features as measured by leg adduction variability and amplitude. These dystonic features were most pronounced during the treadmill task, where mice were forced to continuously ambulate at progressively increasing speeds, as opposed to during open field, where mice were allowed to ambulate at will at their preferred speeds. Compared to term born controls, mice born preterm demonstrated lower foot angle minimums (demonstrating greater leg adduction amplitude) and higher foot angle variance (demonstrating greater leg adduction variability) during treadmill gait (Figure 4A). However, these metrics were not significantly different between mice born preterm and term born controls during open field gait when including both support phases (bipedal and tripedal/quadrupedal support). However, during the less stable bipedal support phase of gait, mice born preterm demonstrated significantly lower foot angle minimums and higher foot angle variances compared to term born mice, suggesting emergency of dystonia during open field when mice were in more challenging periods of the gait cycle (Figure 4B)

**Figure 4.**
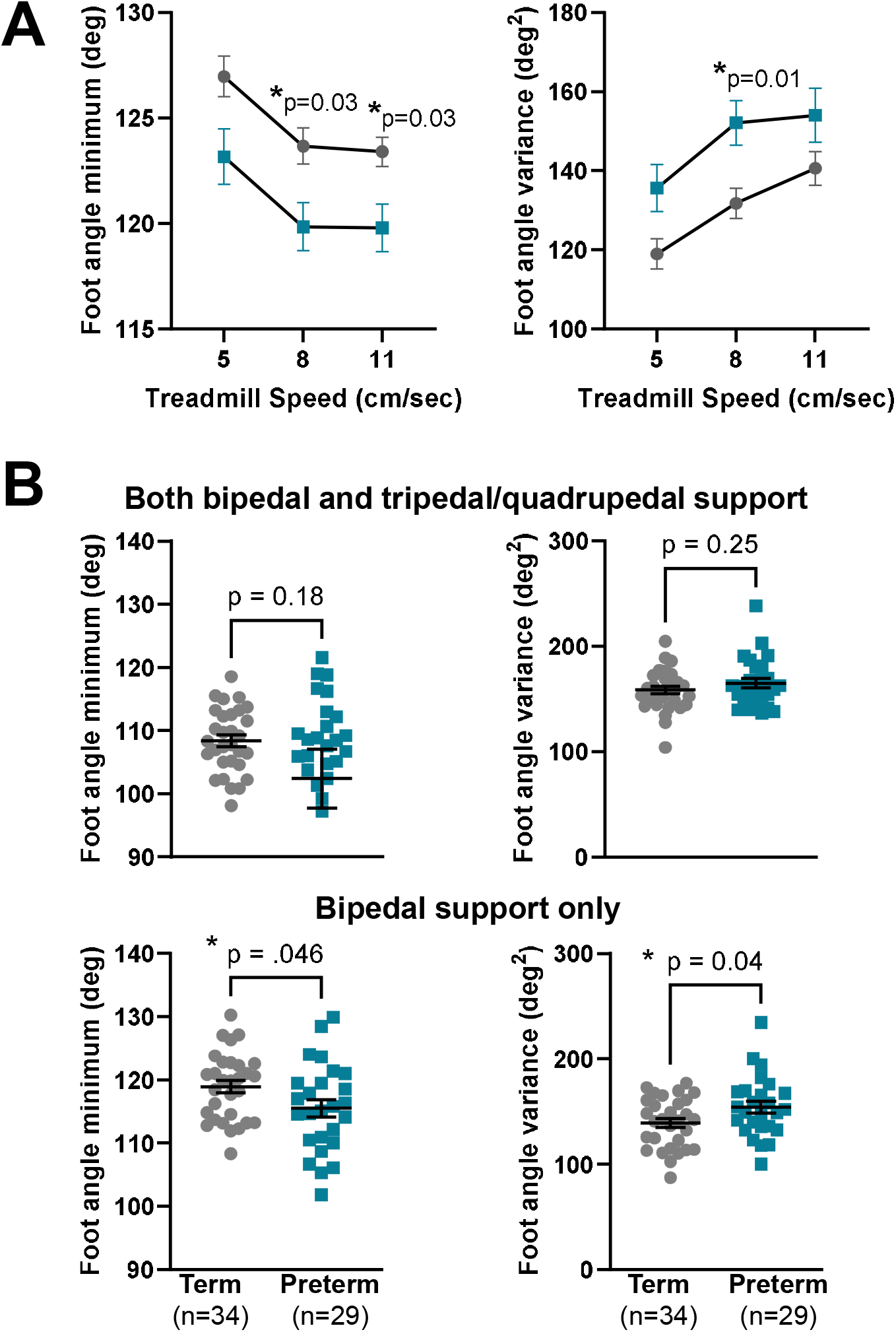
Clinically validated dystonic gait features during treadmill (A, two-way repeated measures ANOVA) and open field (B, T-test). During treadmill gait, mice demonstrate decreased foot angle minimum and variance (A), but during open field, this is only apparent during the less stable bipedal support period of gait.

Finally, to begin assessing cortical and striatal dysfunction in this novel model of preterm birth, we assessed parvalbumin immunoreactivity in the sensorimotor cortex and the striatum (Figure 5). Mice born preterm demonstrated decreased parvalbumin immunoreactivity in the sensorimotor cortex compared to term-born controls as measured by the percent area stained within a standardized sensorimotor cortex ROI (Figure 6A). This percent area difference could encompass both decreased neuropil staining and decreased interneuron number (Figures 5C and 5D). This decrease in parvalbumin immunoreactivity is potentially selective for the cortex: there is no significant difference in parvalbumin neuron number in the striatum between mice born preterm and term-born controls (Figure 6B).

**Figure 5.**
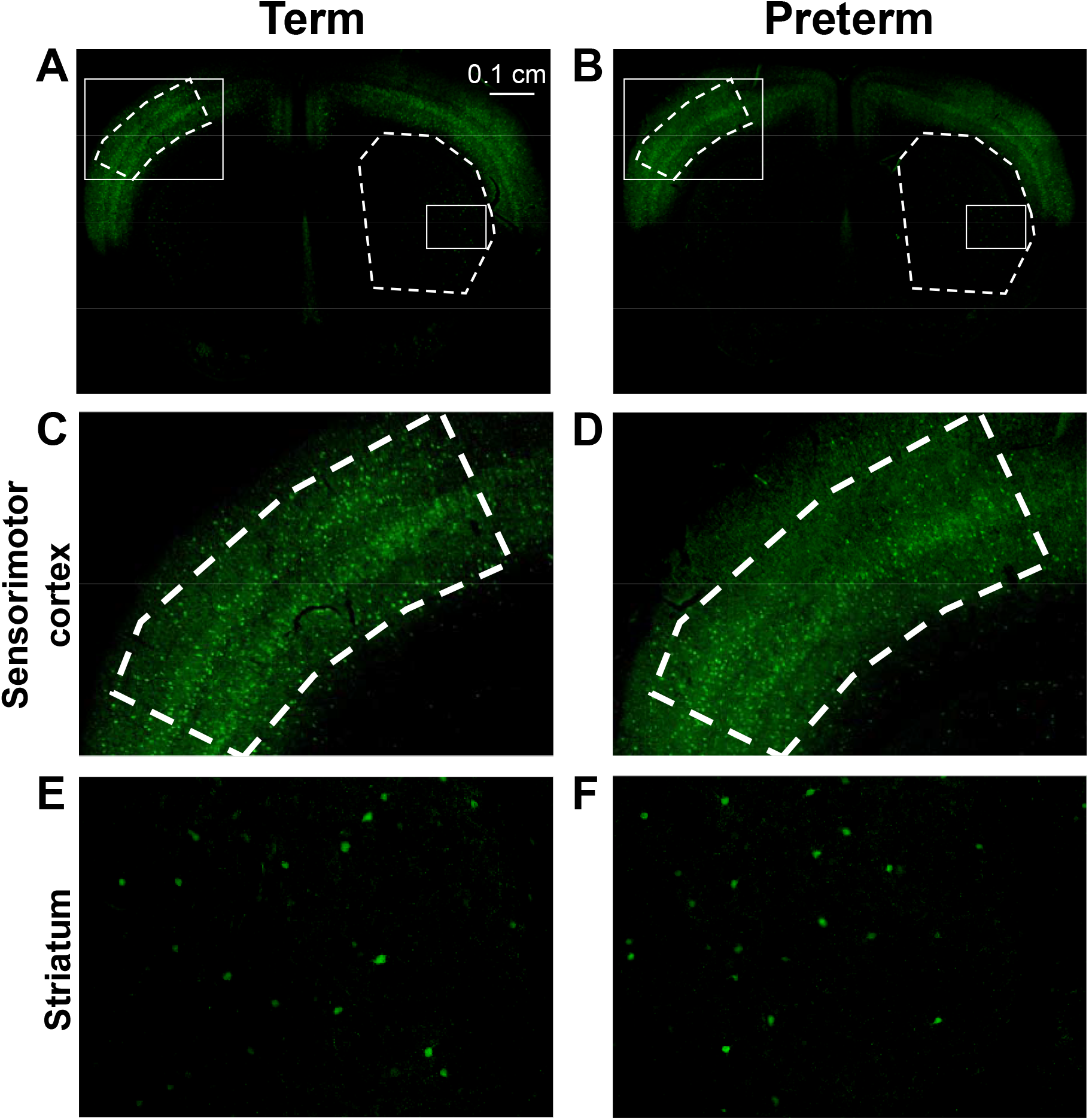
Example parvalbumin immunohistochemistry in mice born at term (A, C, E) and preterm (B, D, E) in an axial brain slice taken at 0.7 mm anterior to bregma (A, B). Regions magnified in C-F are indicated in A and B with rectangular boxes. C,D – sensorimotor cortex. E, F – striatum. Note that there is reduced parvalbumin immunoreactivity in this mouse born preterm compared to a term born control in the sensorimotor cortex (D vs. C) but not in the striatum (F vs. E).

**Figure 6.**
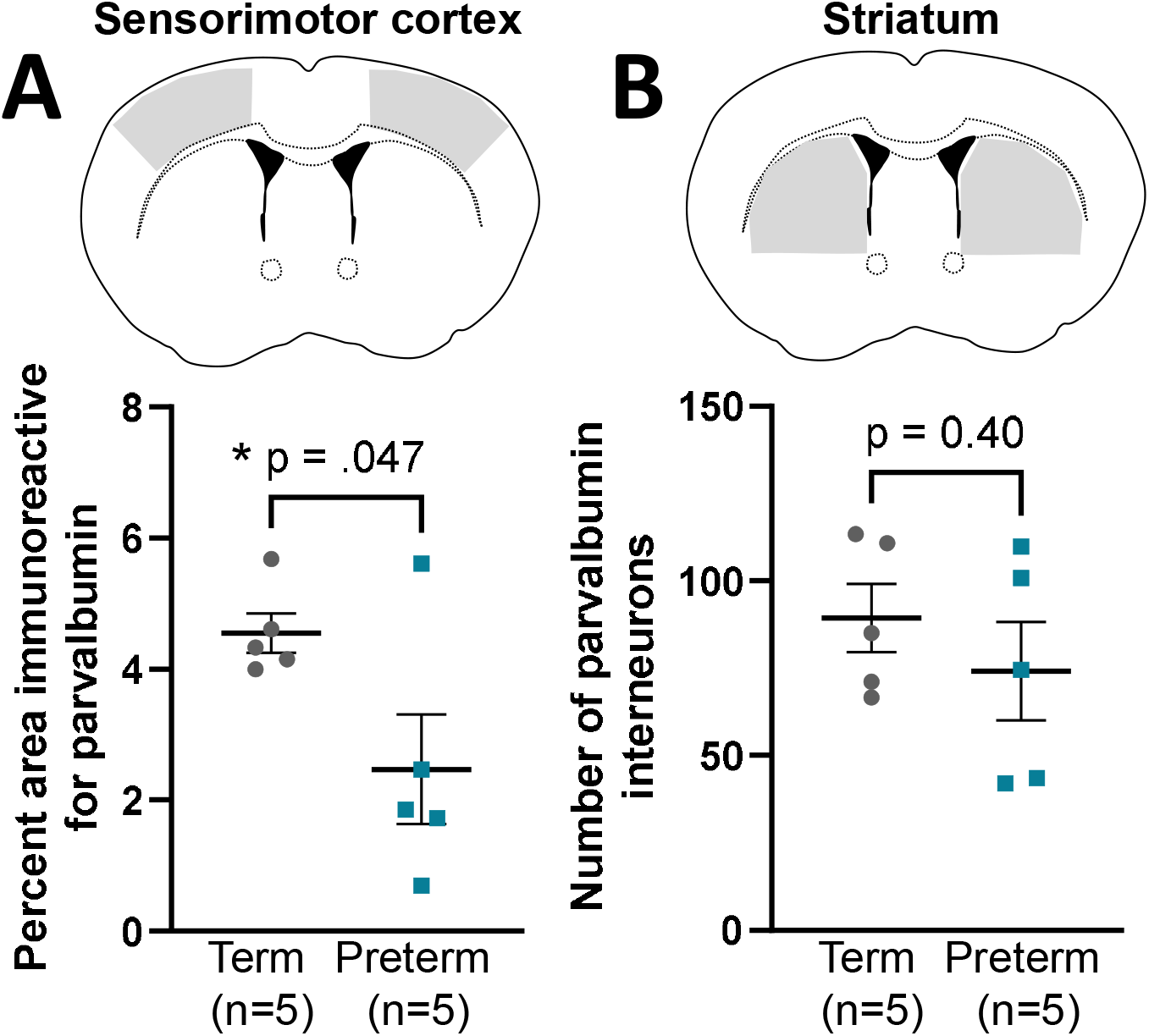
Parvalbumin immunoreactivity in the sensorimotor cortex (A) and striatum (B). Mice born preterm demonstrate reduced parvalbumin immunoreactivity compared to term born controls in the sensorimotor cortex (A), but not in the striatum (B). *T-test

## Discussion

We describe a new mouse model of preterm birth where mice develop subtle locomotor impairment (reduced bipedal support duration) and clinically-validated features of dystonia (leg adduction variability and amplitude). Mice born preterm also have reduced parvalbumin immunoreactivity in the sensorimotor cortex, but not the striatum. These data taken together support clinical data suggesting that abnormal sensorimotor cortex inhibition can cause dystonia following preterm birth. We propose that this model of preterm birth can be used to study potential cortical etiologies and treatment targets of dystonia in CP.

The presence of any locomotor impairment in this model of preterm birth is an improvement upon many existing animal models of dystonia.(Oleas et al., 2013; Richter and Richter, 2014; Tassone et al., 2011) However, the mouse model most commonly used to study CP, the Rice-Vannucci neonatal stroke model, demonstrates much more robust motor impairment and associated cystic or atrophic changes in the cortex and striatum.(Rice et al., 1981; Vannucci and Back, 2022) The histologic findings in mice born preterm are also comparatively subtle, given a lack of obvious cystic or atrophic changes. Future work can also incorporate inflammatory and hypoxic insults together with induction of preterm birth to determine whether that results in a more severe phenotype. However, we note that detailed circuit-based studies to understand dystonia pathophysiology require intact brain tissue to assess. Furthermore, people with CP who are most likely to respond to dystonia interventions clinically are those with largely intact brain tissue such that existing circuits can be modulated with surgical or pharmacologic intervention.(McClelland et al., 2018) Therefore, we view the subtle histologic and locomotor abnormalities demonstrated by this model as a strength for translationally focused dystonia research.

The use of clinically validated dystonia measures in this study also enhances its translational applicability. Clasping, arguably the most extreme form of hindlimb adduction in mice, is primarily elicited during tail suspension, an inarguably difficult task to recapitulate in the clinic. By assessing dystonia during a clinically-relevant task (gait) and also by using clinically-validated measures of dystonia in CP (limb adduction amplitude and variability), it is possible to assess the outcomes assessed in this mouse model directly in the clinic. We propose that our highest likelihood of understanding translationally relevant dystonia pathophysiology comes by assessing clinically validated outcome measures in animal models of disease.(Gemperli et al., 2023)

The role of sensorimotor parvalbumin-positive interneurons in dystonia pathogenesis after preterm birth requires further study. Though the reduced cortical parvalbumin immunoreactivity observed in this study suggests dysfunction of these interneurons, it does not necessarily suggest neuronal loss or reduced neuronal activity. Future work can quantify cortical parvalbumin-positive interneuron number and record from these interneurons *ex vivo* in brain slices prepared from mice born preterm. As we have done with striatal cholinergic interneurons,(Gemperli et al., 2023) direct chemogenetic modulation of sensorimotor parvalbumin-positive interneurons can help determine whether their dysfunction can cause dystonia. Finally, noting that dysfunction of these neurons has also been observed in other conditions that commonly co-exist with CP like autism and epilepsy,(Dos Santos Rufino et al., 2023; Jiang et al., 2016; Juarez and Martínez Cerdeño, 2022; Påhlman et al., 2021) longitudinal characterization of social behavior, cognition, and seizure risk in this model of preterm birth would be valuable.

In sum, we have demonstrated a novel mouse model of preterm birth that demonstrates gait dystonia and evidence of cortical dysfunction. This model can be used to study dystonia following preterm birth and may be useful to study other sequelae of prematurity.

## Author Roles

KG, FF, BN, RJ, CH, RR: methodology, validation, formal analysis, investigation, data curation;

RG: conceptualization, methodology, formal analysis, writing – review and editing;

BA: conceptualization, methodology, validation, formal analysis, resources, data curation, writing – original draft, visualization, supervision, project administration, funding acquisition

## Acknowledgements and Funding

Funding supporting this work is from the National Institutes of Neurological Disorders and Stroke (1K08NS117850-01A1 and 1R01NS112234). There are no conflicts of interest.

